# BAGSE: a Bayesian hierarchical model approach for gene set enrichment analysis

**DOI:** 10.1101/662171

**Authors:** Abhay Hukku, Corbin Quick, Francesca Luca, Roger Pique-Regi, Xiaoquan Wen

## Abstract

Gene set enrichment analysis has been shown to be effective in identifying relevant biological pathways underlying complex diseases. Existing approaches lack the ability to quantify the enrichment levels accurately, hence preventing the enrichment information to be further utilized in both upstream and downstream analyses. A modernized and rigorous approach for gene set enrichment analysis that emphasizes both hypothesis testing and enrichment estimation is much needed. We propose a novel computational method, Bayesian Analysis of Gene Set Enrichment (BAGSE), for gene set enrichment analysis. BAGSE is built on a Bayesian hierarchical model and fully accounts for the uncertainty embedded in the association evidence of individual genes. We adopt an empirical Bayes inference framework to fit the proposed hierarchical model by implementing an efficient EM algorithm. Through simulation studies, we illustrate that BAGSE yields accurate enrichment quantification while achieving similar power as the state-of-the-art methods. Further simulation studies show that BAGSE can effectively utilize the enrichment information to improve the power in gene discovery. Finally, we demonstrate the application of BAGSE in analyzing real data from a differential expression experiment and a Transcriptome-wide Association Study (TWAS). Our results indicate that the proposed statistical framework is effective in aiding the discovery of potentially causal pathways and gene networks. BAGSE is implemented using the C++ programming language and is freely available from https://github.com/xqwen/bagse/. Simulated and real data used in this paper are also available at the Github repository for reproducibility purposes.

## 1 Introduction

Gene set enrichment analysis has become a standard analytic tool in systems biology and bioinformatics. Its primary aim is to identify specific groups of genes in which the association signals are enriched (or depleted) given the association evidence from individual genes. The results from gene set enrichment analysis have implications beyond the association evidence at the single gene level: as the gene set is typically defined by the biological relevance of the member genes, the enrichment of signals in specific gene sets sheds lights on the underlying biological pathways and gene networks, which subsequently helps to uncover relevant molecular mechanisms in a biological system. In practice, gene set enrichment analysis is often conducted downstream of differential expression (DE) analysis and genome-wide genetic association analysis (GWAS). Recently emerged transcriptome-wide association analysis (TWAS) has shown promise in linking causal genes to complex traits utilizing both the data from mapping expression quantitative trait loci (eQTL) and GWAS (Gamazon *et al.*, 2015; Gusev *et al.*, 2018; Zhu *et al.*, 2016). Gene set enrichment analysis based on TWAS results will have the potential to uncover the causal gene networks that lead to complex diseases.

The available gene set enrichment analysis approaches in literature can be roughly classified into two groups. The first group is represented by the popular approach GSEA (Mootha *et al.*, 2003; Subramanian *et al.*, 2005). For each pre-defined gene set, GSEA constructs a ranked list of all member genes based on their association evidence with respect to the phenotype of interest. It then performs a Kolmogorov-Smirnov (KS)-like test to compare the distributions between different gene sets. This procedure has been widely used since its inception, as shown in Keshava Prasad *et al.* (2008); Guttman *et al.* (2009); Shalem *et al.* (2014); Schaub *et al.* (2018). Built upon the algorithm of GSEA, many software packages provide further improvements targeting specific applications (Segrè *et al.*, 2010; Willer *et al.*, 2013; Speliotes *et al.*, 2010). Notably, GSEA-based gene set enrichment analysis has led to breakthroughs in the profiling of cancer cells (Maruschke *et al.*, 2014), and the studies of complex diseases like schizophrenia (Hass *et al.*, 2015) and depression Elovainio *et al.* (2015). A two-stage procedure characterizes the other class of enrichment analysis methods. Taking an example of an enrichment analysis from a DE experiment: in the first stage, genes are classified into either differentially expressed or not based on the association evidence without considering their gene set annotations; in the second stage, a contingency table is constructed according to the DE status and gene set membership of all investigated genes. The resulting contingency table is subsequently used to quantify the enrichment level (by computing the log odds ratio) and testing enrichment (by a chi-squared test or a Fisher’s exact test) for a particular gene set. This method has also been widely applied in the recent literature of genomics and complex disease studies (Richiardi *et al.*, 2015; Walter *et al.*, 2015; Chang *et al.*, 2016).

Despite the popularity of both types of methods in gene set enrichment analysis, they both lack the ability of accurate quantification of enrichment levels for gene sets. The GSEA approach is statistically rigorous in performing hypothesis testing; however, it is not designed to provide an estimation of the enrichment level. The two-stage approach is seemingly intuitive; nevertheless, the classification in the first stage ignores the uncertainty of the gene-level association evidence, which leads to biased estimates of enrichment levels (the details will be explained in Section 2.3). We argue that the accurate quantification of enrichment from a gene set enrichment analysis is critical in many bioinformatics applications. Such information is necessary for comparing the relative importance of multiple gene sets in the same disease or comparing the roles of the same gene set in various conditions.

In this paper, we propose an empirical Bayes procedure, Bayesian Analysis of Gene Set Enrichment (BAGSE), for gene set enrichment analysis. Our computational approach is derived from a hierarchical model. BAGSE is suitable for not only rigorous hypothesis testing but also accurate quantification of enrichment levels. Additionally, BAGSE can simultaneously handle multiple and/or mutually non-exclusive gene set definitions, a feature currently missing from the existing methods. Finally, we show that within the proposed hierarchical model framework of BAGSE, the gene set enrichment information can be subsequently applied for improving the power in uncovering association evidence at the gene-level. The software package implementing the proposed procedures is made freely available at https://github.com/xqwen/bagse/.

## 2 Methods

### 2.1 Model and notations

We consider a general setting suitable for analyzing summary-level data generated from both DE and TWAS studies. Specifically, we use *β*_*i*_ to denote the effect size of the association for each gene *i*. In DE analysis, *β* typically represents the log fold-change of expression levels under two different experimental conditions; in TWAS, *β* quantifies the strength of association between the phenotype of interest and the genotype-predicted gene expression levels. Suppose that the analysis of each gene *i* yields a maximum likelihood estimate of effect size, 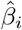, along with its standard error, *ŝ*_*i*_. With sufficient sample size, it follows that

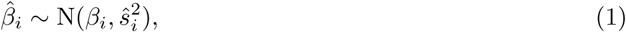

and we also consider that 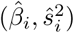 is a sufficient statistic for *β*_*i*_. Throughout this paper, we assume the observed gene-level association data are summarized by 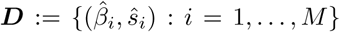 for all *M* genes, and it is made available for enrichment analysis. In the case that only *p*-values are made available, we map each *p*-value to a corresponding *z*-statistic through a standard normal distribution, hence 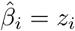 and *ŝ*_*i*_ = 1 Efron (2012); Stephens (2016).

We define a latent binary indicator *γ*_*i*_ to represent the true association status of gene *i*, i.e., *γ*_*i*_ = 1 indicates gene *i* is genuinely associated and 0 otherwise. We assume its annotation data, *d*_*i*_, provides potential prior knowledge on *γ*_*i*_. For the mathematical convenience of the presentation, unless otherwise specified, we assume a single gene set is pre-defined, and *d*_*i*_ is a binary indicator representing if gene *i* is annotated. (In Section 2.4 and the Supplementary Material, we relax this restriction and consider multiple overlapping gene sets.) We assume a logistic prior function connecting *d*_*i*_ and *γ*_*i*_, i.e.,

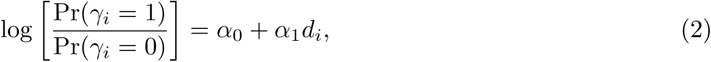

where the coefficients ***α*** := (*α*_0_, *α*_1_) quantify the enrichment information. For example, if *α*_1_ *>* 0, the genes belonging to the gene set of interest are more likely to be associated.

To complete the hierarchical model, we follow the recently proposed adaptive shrinkage (ASH) method (Stephens, 2016) to model the prior effect size *β*_*i*_ (conditional on *γ*_*i*_ = 1) using a mixture of *K* normal distributions, i.e.,

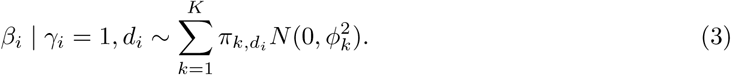

Additionally, conditional on *γ*_*i*_ = 0, *β*_*i*_ = 0 by definition.

In practice, we determine the number of the mixing components, *K*, and corresponding effect size parameters 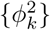 using a data-driven approach as described in Stephens (2016). (The technical details are also described in Section 1.4 of the Supplementary material). Importantly, we allow the mixture proportions, i.e., 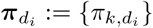, to vary across different types of annotations, which provides the necessary flexibility to model potentially different effect size distributions for different kinds of gene sets or pathways. We view this feature as an improvement and a generalization of the original ASH model.

The proposed Bayesian hierarchical model can be summarized by the graphical model shown in Fig. 1.

**Figure 1:**
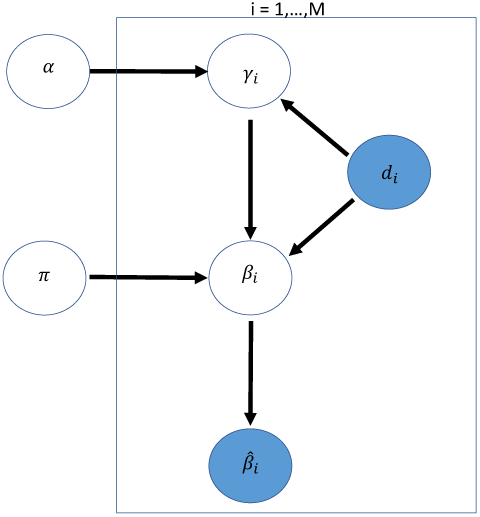
A graphical model representation of the BAGSE model. The directed acyclic graph (DAG) represents a probabilistic generative model. The shaded variables represent data that are observed.

With observed association data ***D*** and annotation data ***d*** := (*d*_1_, …, *d*_*M*_), we frame the problem of enrichment analysis as an inference problem with respect to the enrichment parameter ***α***.

### 2.2 Gene set enrichment estimation

To quantify the enrichment level of annotated genes within a pre-defined gene set, we perform maximum likelihood estimation with respect to ***α*** based on the proposed hierarchical model. Particularly, we design an Expectation and Maximization (EM) algorithm to obtain the maximum likelihood estimates (MLEs) for hyperparameters ***α***, *π* by treating the latent binary vector ***γ*** as missing data.

Briefly, in the E-step of the *t*-th iteration we evaluate the probability 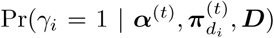 for all genes (where ***α***^(*t*)^ and 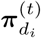 denote the current estimates of ***α*** and 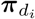, respectively). In this process, the unknown effect size parameters, *β*_*i*_’s, are analytically integrated out. In the M-step, we simply fit a logistic regression model

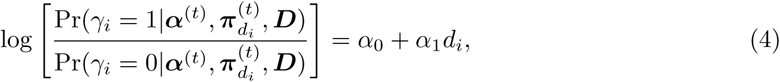

and use the resulting Generalized linear model (GLM) estimates of *α*_0_ and *α*_1_ to obtain the updated ***α***^(*t*+1)^. Subsequently, 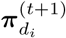 is computed by maximizing a simple *K*-dimensional multinomial likelihood function.

We start the algorithm from a set of arbitrary values of ***α*** and 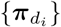 and iterate between the E and M steps until the pre-defined convergence criteria are met. The standard error of 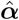 is computed using a profile likelihood approach. The full details of the EM algorithm are provided in the Supplementary Material. Finally, we summarize the result of the gene set enrichment analysis by constructing a 95% confidence interval of *α*_1_ from the EM output. Furthermore, we can obtain a *p*-value by computing a *z* statistic from 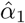 and its standard error to test the null hypothesis,

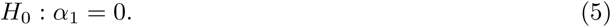

### 2.3 A latent contingency table interpretation

Here we provide an intuitive and general view of our enrichment analysis model and algorithm. Without loss of generality, we consider a single binary annotation for a particular gene set definition (i.e., a gene is either in or out of the annotated pathway/gene set). Now consider an ideal (but unrealistic) scenario where the true association status of each gene is indeed known. Under this setting, the enrichment analysis can be formulated as a 2 × 2 contingency table with the 4 cells indicating the 4 possible combinations of association and annotation status. Given the table, it is straightforward to compute the odds ratio to quantify the level of enrichment. It should be known that the enrichment computation from the contingency table is also statistically equivalent to fitting a simple logistic regression model. However in practice, the exact classification of the association status is unknown, and the above simple procedure is not directly applicable.

As mentioned in the introduction, the commonly applied two-stage procedure can be viewed as an ad-hoc procedure to fill in the unobserved 2 × 2 contingency table based on a simple classification rule. That is, a gene is classified as “associated” if the null hypothesis from a statistical test is rejected, and “unassociated” otherwise. This procedure is intuitive but has some notable caveats. Importantly, it should be clear that the “filled” contingency table is not necessarily accurate compared to the underlying true table. This is because adopting hypothesis testing as a classification procedure preferentially restricts type I errors (false positives) but not the overall classification errors, in which type II (false negatives) errors make up a substantial proportion and are not controlled.

To demonstrate this point, we perform simulations and apply the two-stage procedure to estimate the enrichment parameter. In each simulation, we first generate a 2 × 2 contingency table given the true association and annotation status. We subsequently adjust the cell counts to reflect both the type I and type II errors in the gene-level hypothesis testing. Finally, we compute the enrichment parameter *α*_1_ using the adjusted contingency table. In summary, we find that the two-stage procedure consistently yields the enrichment estimates that are biased toward 0. With the type I error under control, the degree of bias is negatively correlated with the power of gene-level tests. An example from the simulation study is shown in Supplementary Figure 2. Interestingly, the lower power of the gene-level tests is also associated with a higher degree of variation in the enrichment estimates from the adjusted contingency table. Therefore, we conclude that the two-stage procedure can be inaccurate in estimating the enrichment parameter. Nevertheless, for enrichment testing, the direction of the bias from the two-stage procedure seems only to impact power but does not inflate the type I error that asserts enrichment when *α*_1_ = 0, as can be noted from results in Supplementary Table 2.

Our proposed EM algorithm, in this case, can be viewed as an iterative approach to fill in the unobserved contingency table, accounting for the uncertainty of the true binary association status. In the E-step of the proposed EM algorithm, we essentially fill in the table with the expected values of each gene. Note that

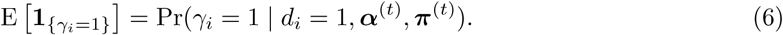

Hence, gene *i* contributes to the cell count of associated and annotated by an amount of Pr(*γ*_*i*_ = 1 | *d*_*i*_ = 1, ***α***^(*t*)^, ***π***^(*t*)^), and to the cell count of unassociated and annotated by an amount of 1 −Pr(*γ*_*i*_ = 1 | *d*_*i*_ = 1, ***α***^(*t*)^, ***π***^(*t*)^). An obvious advantage of this approach is that the uncertainty of the association analysis result is accounted for. The M-step is essentially the same statistical procedure given the contingency table is filled with the expected cell counts.

### 2.4 Local FDR control accounting for gene set enrichment

Another unique advantage of BAGSE is that the estimated enrichment information can be subsequently utilized in identifying truly associated candidate genes. Intuitively, accounting for the quantitative enrichment information boosts the power in identifying signals. This is accomplished by expanding a parametric empirical Bayes framework of local false discovery rate control procedure described in Stephens (2016). Specifically, we consider testing the null hypothesis *H*_0_ : *γ*_*i*_ = 0 for all genes using the enrichment information. For each test, we evaluate the local FDR (lfdr) by plugging in the enrichment estimate 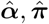,

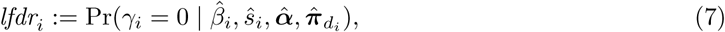

which is a byproduct of our EM algorithm and can be directly applied to control FDR.

Our proposed procedure also provides a principled solution to deal with non-exchangeable multiple hypothesis testing. For example, one may suspect that genes in certain annotated pathways are more likely to be true signals. Such prior expectations can be precisely expressed by equation (2) in our model. In particular, if *α*_1_ > 0 and gene *i* is a member of the gene set of interest, then it has a higher prior probability to be a genuine signal compared to its counterparts absent from the gene set. Furthermore, the enrichment estimation procedure outlined in the EM algorithm allows us to effectively learn the value of *α*_1_ from the observed data. The local FDR computed in (7) combines the enrichment prior and the likelihood information observed from data: if a gene set is estimated to be enriched, the lfdr’s of its member genes are down-weighted by the priors.

In addition to controlling FDR for testing presence or absence of signals (i.e., *γ*_*i*_’s), our approach can be extended to control the local false sign rates (lsfr) (Stephens, 2016), which focus on the signals whose effects can be identified robustly. In particular, we compute lsfr for gene *i* by

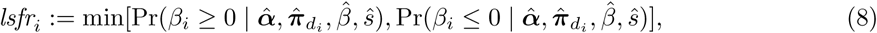

which is interpreted as the error probability in determining the sign of the effect for gene *i*.

### 2.5 Use of multi-category gene set annotations

A unique advantage of BAGSE for enrichment analysis is that it can easily handle multiple gene set annotations, which generalizes the commonly used binary annotations. Consider *L* potentially overlapping gene sets that we wish to evaluate simultaneously. A gene can be independently annotated (i.e., in or out) by an individual annotation. The joint annotation of a gene can be represented by a binary *L*-vector, which can take 2^*L*^ possible values: e.g., (0, 0, …0) indicates a gene is not annotated in any gene set, and (1, 1, …1) indicates a gene is annotated in all gene sets. Note that our parametric prior model (2) can accommodate such multi-category annotation by regarding the corresponding *d*_*i*_ as a categorical variable coded by dummy variables (Section 1.2 of the Supplementary Material). For inference, instead of reporting a single enrichment coefficient (*α*_1_), use of multi-category annotation leads to up to 2^*L*^ −1 enrichment coefficients for different annotation configurations in contrast to the baseline annotation (0, 0, …, 0).

The general scheme applies to an arbitrary number of gene sets (*L*) considered, although a large number of *L* values may lead to expensive computational costs. It is possible to make simplifying assumptions to reduce computational complexity. For example, let binary indicator *u*_*i,l*_ denote if gene *i* is annotated in the gene set *l*. We consider an additive prior model

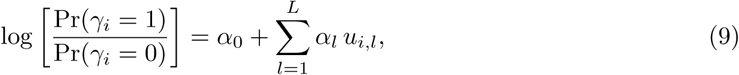

which is computationally feasible for moderate to large *L* values. This particular prior model is also implemented in the BAGSE.

One of the unique advantages of the multi-category gene set annotation is to maximize the utility of the proposed FDR control procedure. In GO and KEGG pathways, a single gene is often annotated in multiple gene sets. Relying on a single gene set to determine the local FDR of the given gene seems sub-optimal. The logical alternative is to combine the multiple gene sets and generate a single multi-category gene set annotation. We argue that such annotation provides objective and optimal FDR control to maximize the discovery power by incorporating information from gene set annotations. We demonstrate this point in Section 3.3.

## 3 Results

### 3.1 Simulation studies

We use numerical simulations to benchmark the performance of the proposed Bayesian gene-set enrichment analysis procedure. Our particular focuses are put on examining the accuracy of the enrichment estimates and its performance in enrichment testing.

In each simulated dataset, we consider the analysis of 10,000 genes, each of which is assigned a binary true association status based on the enrichment parameters (*α*_0_, *α*_1_) and pre-defined annotations. For each gene, we assume an association *z*-score is available for the enrichment analysis: for un-associated genes, we draw the *z*-scores from the standard normal distribution; for the associated genes, the *z*-scores are simulated from a *t*-distribution with the degree of freedom = 10 (which mimics the long-tailed effect size distribution commonly observed in practice, see Supplementary Figure 1 for example). We set the enrichment parameter *α*_0_ = −1 throughout, while varying the values of *α*_1_ and the proportion of annotated genes (denoted by *q*) across all simulations. We vary *α*_1_ values from 0 to 1, due to all investigated methods having power similarly near 100% power as *α*_1_ increases to above 1. We use values for *q* ranging from 1% to 20%, due to that being a realistic range for *q* based on the hierarchical nature of popular databases such as KEGG or GO. Each parameter set is used to generate 5000 different datasets.

#### 3.1.1. Evaluation of enrichment parameter estimation

We first examine the point estimate of the enrichment parameter from the BAGSE analysis procedure. For comparison, we also compute the enrichment estimate from the two-stage procedure. The analysis results of the simulated datasets are summarized in Fig. 2 for annotation proportion *q* = 10%. It is clear that the proposed approach consistently yields unbiased enrichment estimates across all *α*_1_ values. Also consistent with the previous results, the estimates from the two-stage approach are biased towards 0. Across different *q* values, the estimates from the Bayesian procedure remain unbiased. But we observe a pattern that shows a lower level of variation in 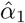 when *q* increases towards 0.5. The latent contingency table interpretation can intuitively explain this phenomenon: as *q* tends to 0 (or 1), the underlying table becomes more imbalanced, causing increased uncertainty for the enrichment parameter. Results from simulations using all parameter sets can be found in Supplementary Table 1 and Supplementary Figures 3, 4 and 5.

**Figure 2:**
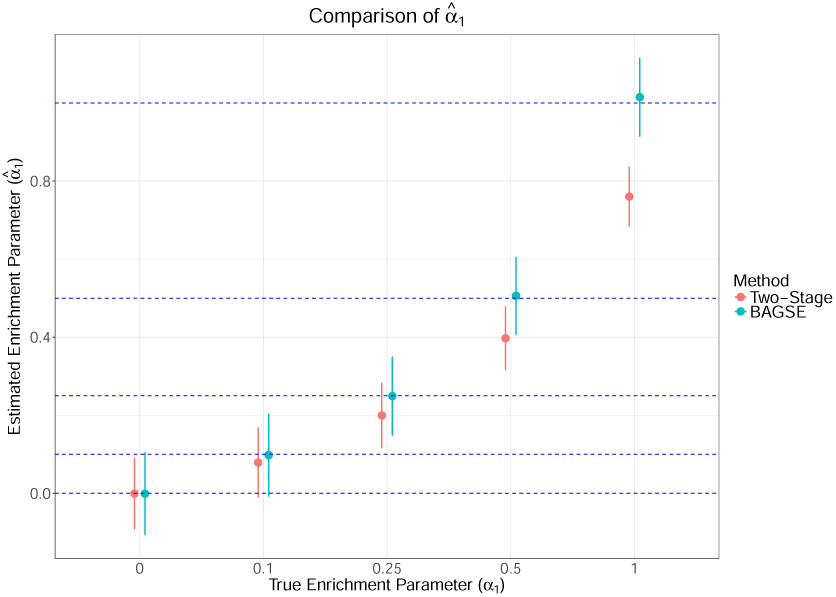
A comparison of the enrichment parameter estimates between the two-stage approach and the proposed method, with standard errors represented as error bars. BAGSE appears to give an unbiased estimate of *α*_1_, while the two-stage approach’s estimate grows more severely biased as the enrichment parameter grows higher.

We conduct additional simulations to illustrate the ability of the proposed method in estimating enrichment parameters using annotations from multiple overlapping gene sets. In summary, we find that BAGSE estimates in such scenario remain unbiased and accurate. The details of these additional simulations are described in Section 2 of the Supplementary Material.

#### 3.1.2. Power comparison in enrichment testing

Next, we proceed to examine the performance of various methods, including BAGSE, the two-stage approach, and the popular GSEA method, in testing the enrichment hypothesis: *H*_0_ : *α*_1_ = 0. We apply both the unweighted and weighted forms of the GSEA procedure. The unweighted GSEA procedure corresponds to a standard Kolmogorov-Smirnov test (by comparing the distribution of the absolute value of *z*-scores, or *p*-values, between annotated and unannotated genes). We use the default weight recommended in the original GSEA paper Subramanian *et al.* (2005) to perform the weighted GSEA procedure.

The simulation results for when *q* = 10% are summarized in Fig. 3. We conclude that all methods properly control type I errors at 5% level, based on the proportion of tests rejected when *α*_1_ = 0 being below 0.05 (denoted by the dotted line) for all methods. BAGSE and the weighted GSEA method are top performers at every combination of simulation parameters, constantly outperforming the unweighted GSEA and the two-stage approaches by a significant margin, especially at the intermediate *α*_1_ values of 0.1 and 0.25. The power difference between BAGSE and the weighted GSEA is generally negligible. In our simulation setting, when *α*_1_ → 1, all methods seemingly achieve the perfect power to reject the null hypothesis. Additionally, we observe that the power to detect enrichment improves as *q* increases towards 0.5. This phenomenon can be similarly explained by the latent contingency table interpretation of our proposed model: the standard error of the enrichment estimate decreases as the proportion of the annotation increase towards 0.5. Results from simulations using all parameter sets for all methods can be found in Supplementary Table 2 and Supplementary Figures 6, 7 and 8.

**Figure 3:**
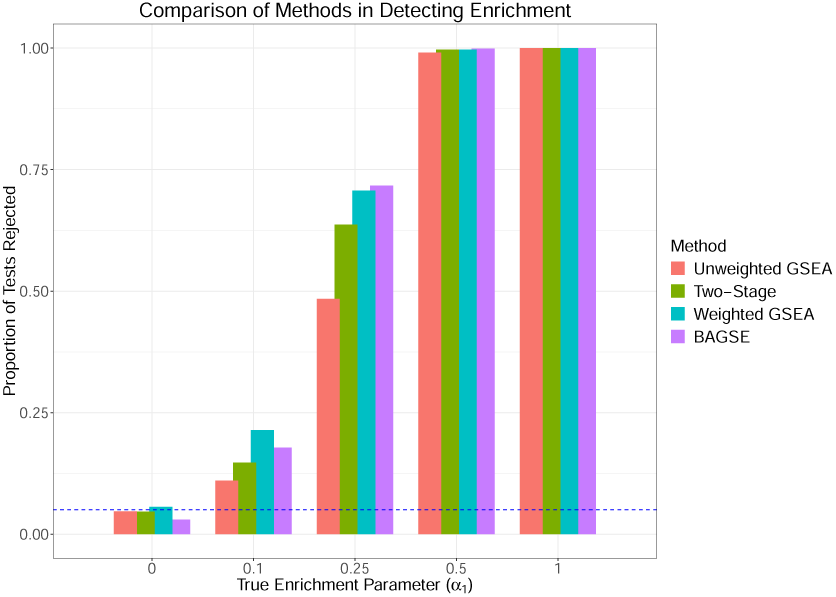
A comparison of type I error and power to detect gene set enrichment using unweighted GSEA, two-stage approach, weighted GSEA, and BAGSE. The *y*-axis denote the proportions of simulated data sets where the null hypothesis are rejected The leftmost columns, corresponding to the null model *H*_0_ : *α*_1_ = 0, represent the type I error rates for the methods. As the enrichment parameter takes non-zero values, the data are simulated under the alternative scenarios, and the corresponding columns represent the power. All methods control the type I errors properly. As expected, power for all methods increases as the true enrichment parameter increases. BAGSE and weighted GSEA outperform the other methods in this regard.

#### 3.1.3 Gene discovery incorporating enrichment quantification

Finally, we examine the power in identifying truly associated genes when gene set enrichment information is explicitly considered. Specifically, we compute the local FDR for each gene using (7) where the estimated enrichment parameter is plugged in, and control the overall FDR at 5% level. For a baseline comparison, we use both the *q*-value procedure Storey *et al.* (2003) and the local FDR procedure (both are implemented in the R package “*qvalue*”) to control FDR at the same level ignoring the gene set information.

Our results indicate that all methods properly control the FDR at the desired level. However, the proposed procedure accounting for enrichment quantification consistently outperforms the *q*-value and the local FDR procedure ignoring the enrichment information in realized power (Figure 4). As expected, the improvement of power is positively correlated with the level of enrichment. In our simulation setting, we observe a modest increase in power (∼ 1.5%, or 20 more true positive discoveries per simulated dataset) when the enrichment parameter is ∼ 1; as *α*_1_ reaches 5, the power boost becomes much more substantial (∼20% improvement in power, or 400 more true discoveries per simulated dataset).

**Figure 4:**
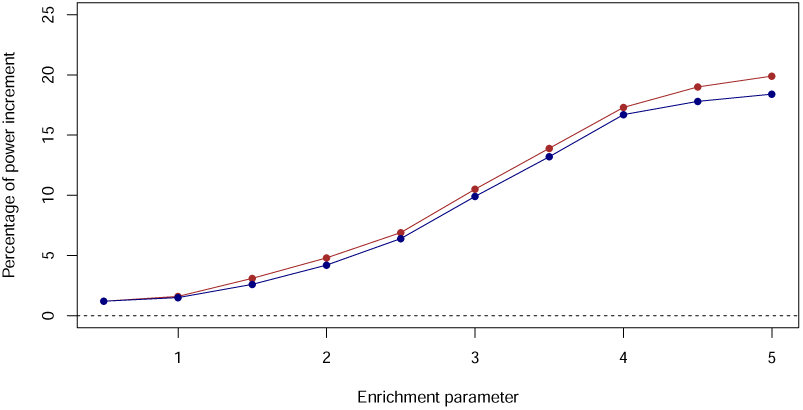
Power increment in gene discovery when accounting for gene set enrichment information. The percentages of power increment by using enrichment information are compared to two standard FDR control procedures: the *q*-value method (brown line) and the local FDR method (blue line). All methods control FDR at 5% level.

### 3.2 Real data application I: differential expression experiment

Next, we apply BAGSE to the experimental data from Moyerbrailean *et al.* (2016), which considers the differential expression of genes under the treatment of glucocorticoid. The study profiles 20,896 genes using RNA-seq. A *p*-value is obtained for each gene using the software package DESeq2 (Love *et al.*, 2014). We convert the p-values into the z-scores, using the estimated effect sizes from the study to determine the corresponding signs. The experiments are also carried out in multiple tissues. For demonstration purposes, we select the results gathered from the peripheral blood mononuclear cells. Because of the known nature of these cells in their response to glucocorticoid, we expect genes involved in pathways associated with immune response to be enriched.

There are 21 KEGG pathways involved in the immune responses. Of these, 15 contain genes that are in our dataset. We first analyze these 15 pathways separately using BAGSE. The enrichment estimate for each of these pathways is provided in Supplementary Table 3. We observe that the majority of the examined pathways (12 out of 15) are shown to be significantly enriched with DE genes, and their enrichment levels can be straightforwardly compared by utilizing the quantified enrichment estimates. In particular, we find that the Intestinal immune Network for IgA Production pathway seemingly shows an extremely high level of enrichment (*α*_1_ = 9.34 with 95% CI [5.38, 13.29]).

Because each pathway only annotates a small proportion of genes, the confidence intervals are typically large (Supplementary Table 3). Following Carbonetto and Stephens (2013), we pool the genes annotated in the 15 KEGG pathways to form a general category of the gene set and examine the enrichment of DE genes in this gene set representing general immune responses. In total, 2.7% of the 20,896 genes are annotated in the aggregated gene set.

We apply BAGSE, the two-stage approach, and GSEA with the two different weighting schemes to conduct gene set enrichment analysis. These results are summarized in Table 1. BAGSE detected strong enrichment for the pooled immune response gene sets, with an enrichment log odds ratio of 1.31 (95% CI [0.96,1.66]), which corresponds to a *p*-value of 2.2 × 10^−13^. The two-stage approach also detects enrichment, with an enrichment log odds ratio of 0.87 (95% CI [0.63, 1.11]). The unweighted GSEA method detects significant enrichment with a p-value of 1 × 10^−7^. The weighted GSEA method also detects significant enrichment with an estimated p-value *<* 1 × 10^−3^.

**Table 1:**
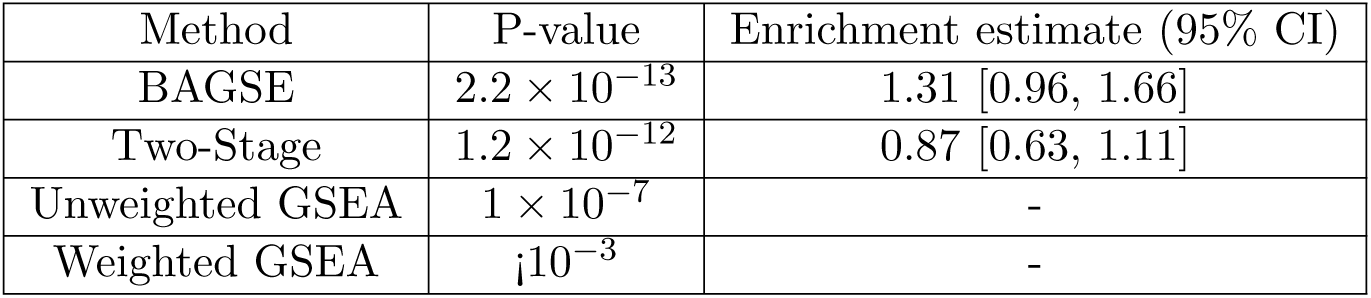
The comparison of significance in detecting enrichment between methods for data from the Differential Expression of genes under treatment of glucocorticoid. As expected, all methods show significance. Note that the inexact value for the weighted GSEA p-value is due to its calculation using 1,000 permutations (recommended by Subramanian *et al.* (2005)).

Additionally, we examine the power improvement in identifying DE genes by incorporating the quantified enrichment information. Without using the enrichment information, we find that the *q*-value procedure identifies 1496 genes at 5% FDR level (the local FDR procedure, implemented in the *q*-value package, identifies 1527 genes). By incorporating the enrichment estimates, BAGSE identifies 1617 genes at the same FDR control level, which amounts to 8% increase in comparison to the *q*-value procedure (or 6% increase to the local FDR procedure). Overall, these results appear consistent with our simulation studies.

### 3.3 Real data application II: transcriptome-wide association analysis

For the second illustration with real data, we perform gene set enrichment analysis using the association results generated from the TWAS analysis. TWAS is a principled approach based on Mendelian randomization (MR) and attempts to test potential causal relationships from the expressions of candidate genes to the complex trait of interest (Gamazon *et al.*, 2015; Gusev *et al.*, 2018; Zhu *et al.*, 2016). Specifically, TWAS analysis constructs a genetic gene expression prediction function using the eQTL data and examines the association between the predicted gene expression levels and the complex trait of interest from the GWAS data.

In this example, we use the GWAS summary statistics of high-density lipoprotein (HDL) levels from the Global Lipids Genetics Consortium (GLGC) (Willer *et al.*, 2013) and the multi-tissue eQTL data from the GTEx project (version 8). We apply the S-PrediXcan (Barbeira *et al.*, 2016) algorithm and obtain TWAS *p*-values for a total of 32,362 genes that are investigated in the GTEx project.

Based on the TWAS association statistics, we investigate the enrichment of six gene sets from KEGG and GO. These gene sets are implicated in the original GLGC analysis Willer *et al.* (2013) By the MAGENTA method Segrè *et al.* (2010). The original analysis uses a proximity-based approach and painstaking biological analysis to link a GWAS hit to a nearby gene. (In comparison, the TWAS analysis establishes the links automatically with the aid of additional eQTL data and few subjective criteria.) With the gene-level quantification directly obtained from the TWAS analysis, we aim to re-examine the results, which have been considered as a gold standard in lipids genetic research.

The results of enrichment analysis of the six gene sets by BAGSE and weighted GSEA are summarized in Table 2. At the significance level of 0.05, BAGSE identifies enrichment in five of the six gene sets, while weighted GSEA identifies enrichment in four. We note that, in the sole gene set (Triglyceride Lipase Activity) that barely misses the significance threshold in our BAGSE analysis, the fewest genes (15) are annotated. Consistently with what we observe in the simulation studies, precisely quantifying enrichment (i.e., with reasonably small standard errors) for such small gene sets is highly challenging. The enrichment estimates also enable direct comparison of the relative importance of these relevant biological pathways. It is not surprising that Cholesterol Metabolic Process has the highest level of enrichment among the gene sets examined. But most importantly, this application illustrates that our proposed enrichment analysis framework has the potential to extend the causal implications derived from the TWAS analysis, and uncover the underlying causal molecular networks.

**Table 2:** The results from applying BAGSE and weighted GSEA to the gene sets implicated in (Willer *et al.*, 2013). All gene sets, with the exception of the Triglyceride Lipase Activity, show enrichment at the 5% significance level according to BAGSE.

| Pathway | BAGSE enrichment estimate (95% CI) | BAGSE p-value | Weighted GSEA p-value |
| --- | --- | --- | --- |
| Neurotrophin Signaling Pathway (KEGG) | 1.016 [0.003, 2.028] | 0.049 | 0.052 |
| Adipocytokine Signaling Pathway (KEGG) | 1.237 [0.197, 2.277] | 0.019 | 0.188 |
| Triglyceride Lipase Activity (GO) | 1.614 [-0.008, 3.235] | 0.051 | 0.001 |
| Cholesterol Metabolic Process (GO) | 2.198 [1.387, 3.009] | $1.1 \times 10^{-7}$ | $< 10^{-3}$ |
| Enzyme Binding (GO) | 1.042 [0.322, 1.762] | $4.5 \times 10^{-3}$ | $< 10^{-3}$ |
| Phospholipid Binding (GO) | 1.569 [0.787, 2.352] | $8.4 \times 10^{-5}$ | $< 10^{-3}$ |

Additionally, we again confirm that incorporating enrichment information helps improve the power of association discovery. In this instance, the original TWAS analysis (without enrichment information) identify 578 unique genes at FDR 5% level. By incorporating the enrichment estimate from Cholesterol Metabolic Process, we identify additional 30 genes at the same FDR level, representing a 5% increase in discovery. Finally, we combine the three significant GO terms into a single multicategory gene set annotation and repeat the enrichment estimation and the FDR control procedure. In the end, we find 626 genes are rejected at the FDR 5% level, representing an 8% increase in discovery.

## 4 Conclusion and discussion

In this paper, we have introduced an empirical Bayes procedure, denoted as BAGSE, for gene set enrichment analysis. We have shown, through simulations and real data analysis, that the proposed approach provides a principled inference procedure to estimate the level of enrichment for a gene set, avoiding the caveats of the two-stage procedure. In addition, the proposed Bayesian method maintains strong power in testing enrichment, as compared to other popular approaches to gene set enrichment analysis. Finally, we show that the enrichment estimates from the proposed approach can be subsequently utilized to improve power for gene-level testing.

It should be noted that our approach can be straightforwardly applied to simultaneously estimate the enrichment of multiple gene sets with overlapping genes. This unique feature can be critical because, as seen in many commonly used gene pathway definitions (e.g., the KEGG pathway database), there are many genes involved in multiple pathways. This flexibility in gene annotation intrinsic to the proposed Bayesian model gives it a distinct advantage as an enrichment analysis method.

During our simulations, we noted that both power to detect enrichment and enrichment estimation were heavily dependent on the proportion of genes annotated. Our latent contingency table interpretation can partially explain this phenomenon: even if all association status is indeed observed, the standard error of the enrichment estimate is known to be negatively correlated with the smallest cell count, and lower annotation proportion tends to decrease the smallest cell count (which typically is the cell corresponds to the count of both annotated and associated genes). All enrichment testing and estimation methods would be affected by a low proportion of annotated genes, reflected by losing either power or precision. In general, the quality of the gene set definition has a direct impact on results from enrichment analysis. Defining highly specific gene pathways is an ongoing challenge.

In our real data applications, we illustrate the proposed gene set enrichment analysis based on TWAS result. TWAS analysis focuses on assessing potential causal relationship from a single gene to a complex trait of interest. The gene set enrichment analysis based on TWAS results has the potential to extend causal implications of single genes and uncover potentially *causal* biological pathways. We expect that this type of analysis will become increasingly popular in the field of systems biology. Our proposed approach has the unique advantage in accurate quantification of relative enrichment levels of candidate gene sets, which should help to compare and select causal molecular pathways of complex diseases.

## Supplementary Materials

### A EM algorithm for parameter estimation

Recall that 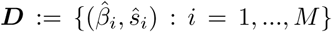 and ***γ*** := (*γ*_1_, …, *γ*_*M*_). We consider the complete data likelihood Pr(***D, γ*** | ***d, α, π***) by treating ***γ*** as missing data:

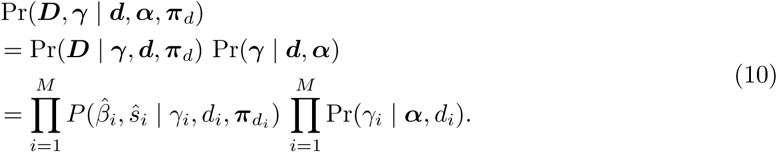

Our model assumes that

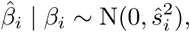

and the prior for *β*_*i*_ follows a *K*-component mixture normal distribution depending on the corresponding gene set annotation, i.e.,

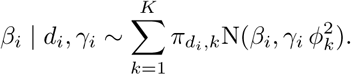

Equivalently, it follows that

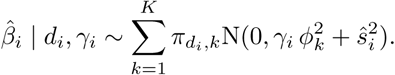

#### A.1 Model reparameterization

We now consider a general case with *L* potentially overlapping gene sets. By the coding convention introduced in section 2.4, the gene set annotation *d*_*i*_ is a categorical variable representing *R* multiple mutually exclusive categories, where 2 ≤ *R* ≤ 2^*L*^. Specifically, we use *d*_*i*_ = 0 to denote the baseline level that a gene is not annotated in any of the *L* gene sets. Under this formulation, the equation (2) in the main text is generalized to

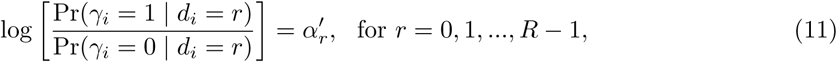

in this coding system. In case that *L* = 1, this coding system is compatible to equation (2). In particular, 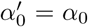 and 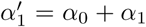. Given *K* pre-defined mixture components for the effect size distribution in the alternative models, ***π*** is represented by a *R* × *K* matrix with (*r* + 1)-th row vector representing the weights for annotation *d*_*i*_ = *r* (*r* = 0, …, *R* − 1).

For each gene *i*, we introduce a [*R*(*K* + 1)]-dimension latent indicator, ***η***_*i*_, to reparametrize the complete data likelihood function (10). Note that all entries of a valid ***η***_*i*_ vector take value 0 except that a single entry takes value 1. In particular, we define

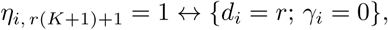

for *r* = 0, …, *R* − 1, and

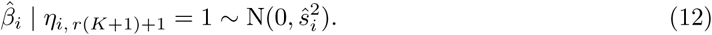

That is, this subset of latent indicators and the corresponding parameters define the null (i.e., non-association) models for each annotation category *r*. Similarly, we set

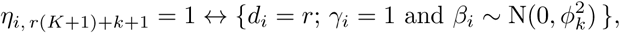

for *k* = 1, …, *K*, and *r* = 0, …, *R* − 1. Accordingly,

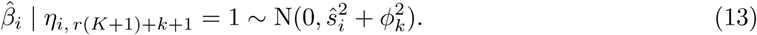

Furthermore, the prior weights for the indicator ***η***_*i*_ are simple functions of the original parameters (***α, π***), i.e.,

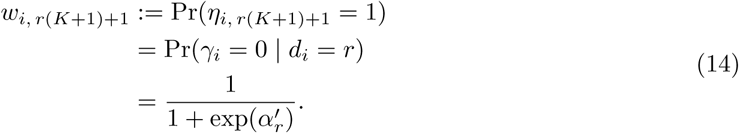

Similarly,

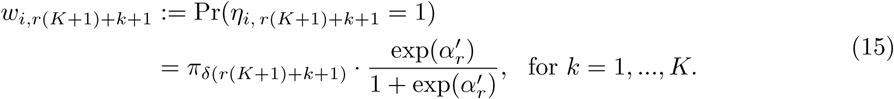

Note that we use the function *δ*(*j*) = (*d*_*i*_ = *r, k*) to map the single index *j* = *r*(*K* + 1) + *k* + 1 to the double indcies *d*_*i*_ = *r* and *k* in the original (***α, π***) notation for suitable integer *j* ∈ *S*, where S denote the index set excluding *j* = *r*(*K* + 1) + 1, for *r* = 0, …, *R* − 1. Similarly, we use *δ*^−1^(*d*_*i*_ = *r, k*) = *r*(*K* + 1) + *k* + 1 to denote the inverse mapping.

Under this alternative parameterization, the complete data likelihood can be expressed by

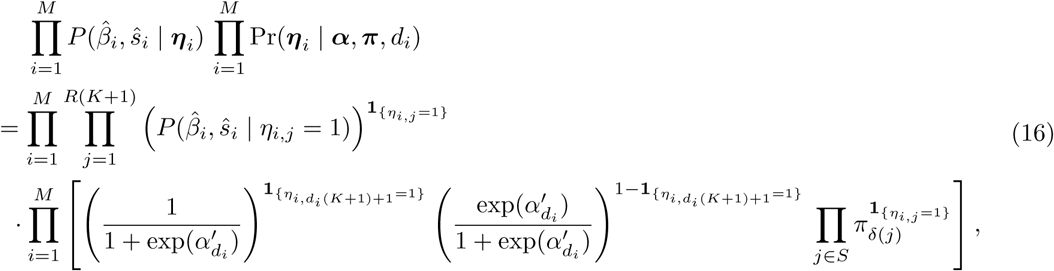

Thus, the complete data log-likelihood is given by

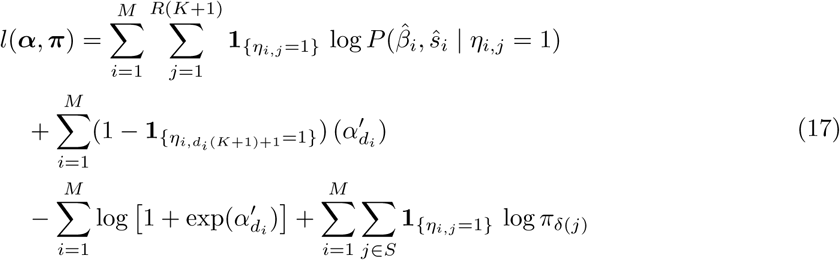

#### A.2 E-step

In the E-step for the *t*-th iteration of the EM algorithm, we compute the expected value of the missing data, ***η***_*i*_, conditional on the observed data summary statistics, gene set annotation, and the current estimates of the parameters, ***α***^(*t*)^, ***π***^(*t*)^, i.e.,

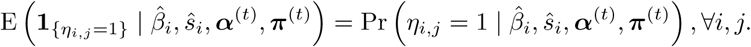

The computation can be carried out by the Bayes rule, i.e.,

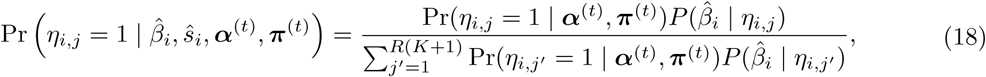

where the relevant likelihood and prior functions are given by (12), (13), (14), and (15), respectively.

#### A.3 M-step

In the M-step for the *t*-th iteration, we update the estimate of the enrichment parameter by finding

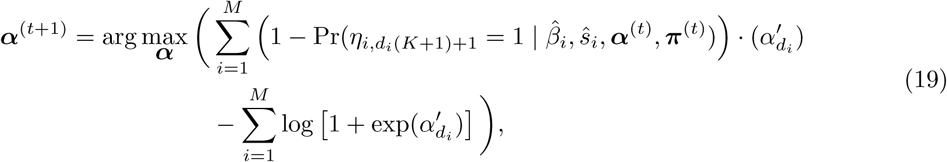

which is equivalent to fitting a logistic regression model with a single categorical covariate, ***d***, and the binary response variables, *η*’s, replaced by their corresponding posterior probabilities. Our implementation uses the Newton-Raphson algorithm to perform numerical optimization. Similarly, we update the estimate of ***π*** by finding

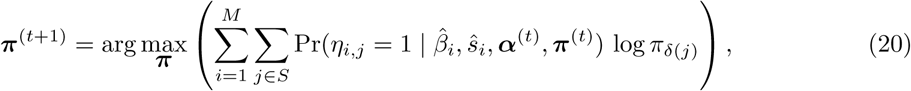

under the constraint

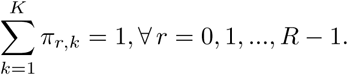

Note that maximization can be achieved analytically. Specifically,

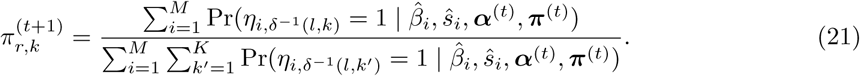

The software implementation of the EM algorithm can initialize the parameters of interest at arbitrary starting points. By default (i.e., without explicit user specification), we set

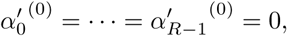

and

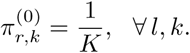

We iterate between the E-step and the M-step until pre-defined convergence criteria is satisfied.

#### A.4 Grid construction

We follow the procedure described in Stephens (2016) to construct a dense set of 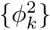 in a data-driven fashion. Specifically, we construct a series of *ϕ*_*k*_ values within the range of (*ϕ*_min_, *ϕ*_max_), where *ϕ*_min_ = min(*ŝ*_*i*_)*/*10 and 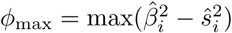. We start with *ϕ*_1_ = *ϕ*_min_ and set

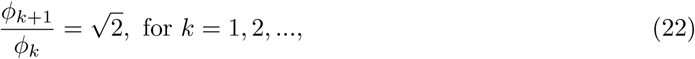

until *ϕ*_*K*+1_ *> ϕ*_max_.

### B Simulation with annotations from multiple gene sets

We perform additional simulations to illustrate the utility of estimating enrichment parameters from multiple overlapping gene sets.

As a proof of concept, we simulate two sets of independent gene set annotations for 10,000 genes in each simulated data set where *q* ≈ 40% of genes are independently annotated in each gene set. Let binary indicators *u*_*i*,1_ and *u*_*i*,2_ denote the annotation status of gene *i* in the two gene sets, respectively. As a result, 36% genes are expected to be not annotated for any of the gene sets (denoted by [*u*_*i*,1_ = 0, *u*_*i*,2_ = 0] or *d*_*i*_ = 0); 24% genes are expected to be annotated only for the first gene set (denoted by [*u*_*i*,1_ = 1, *u*_*i*,2_ = 0] or *d*_*i*_ = 1); 24% genes are expected to be annotated only for the second gene set (denoted by [*u*_*i*,1_ = 0, *u*_*i*,2_ = 1] or *d*_*i*_ = 2); and 16% genes are expected to be annotated for both gene sets (denoted by [*u*_*i*,1_ = 1, *u*_*i*,2_ = 1] or *d*_*i*_ = 3).

For each gene *i*, we draw its association status *γ*_*i*_ from a Bernoulli(*p*_*i*_) distribution, where

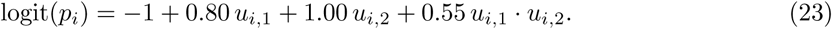

Conditional on *γ*_*i*_ = 1, the corresponding *z* score is drawn from a *t*-distribution with 10 degree of freedom for all *d*_*i*_ values. Otherwise, when *γ*_*i*_ = 0, the *z* score is drawn from the standard normal distribution. We generate 1,000 simulated data sets using the above scheme.

When analyzing the simulated data, we do not assume any knowledge of (23) and simply apply the general combinatorial annotation model and the EM algorithm described in section 2.4 of the main text and section 1 of the supplementary material. Specifically, we examine the enrichment estimates in contrast to the baseline annotation (annotation “00”), i.e., the estimates of 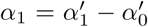 (truth = 0.80) for genes annotated only in set 1 (annotation “10”), 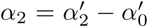 (truth = 1.00) for genes annotated only in set 2 (annotation “01”), and 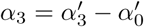 (truth = 0.80 + 1.00 + 0.55 = 2.35) for genes annotated in both set 1 and set 2 (annotation “11”), respectively. Across 1,000 simulations, we find that the proposed EM algorithm yields unbiased estimates of enrichment parameters (Figure 5). We find that all three estimates are seemingly unbiased: the average point estimates for the three mutually exclusive annotations are 0.806, 1.016 and 2.377, respectively.

**Figure 5:**
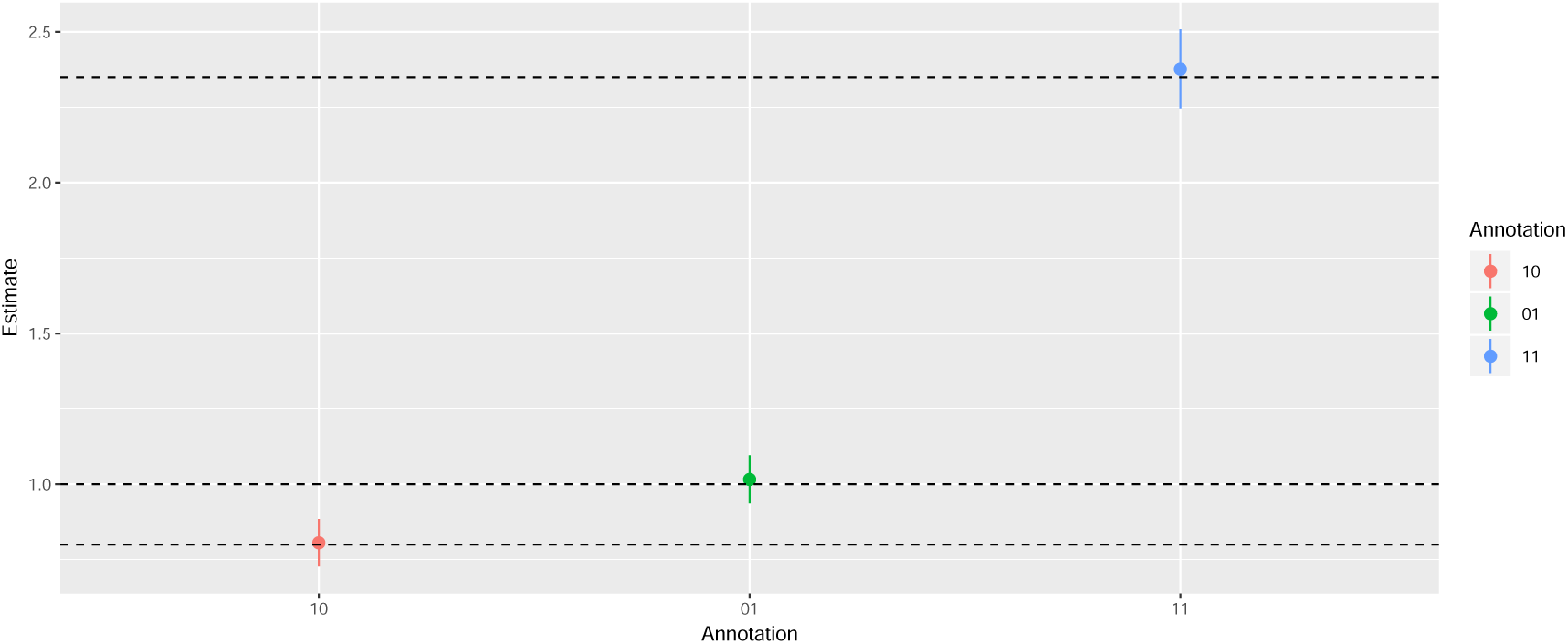
Simultaneous enrichment estimation of overlapping gene sets by BAGSE in simulation studies. Two overlapping gene sets are used in the simulation. Annotation “00” denotes the genes not annotated in any gene set, these genes are the baseline for the enrichment comparison. Annotation “10” denotes the genes annotated only in the first gene set, annotation “01” denotes the genes annotated only in the second gene set, and annotation “11” denotes the genes annotated in both gene sets. The average point estimates and the corresponding standard errors from 1000 independent simulated data sets are plotted. The dashed lines denote the underlying true enrichment levels for each group.

## References

Barbeira, A., Shah, K. P., Torres, J. M., Wheeler, H. E., Torstenson, E. S., Edwards, T., Garcia, T., Bell, G. I., Nicolae, D., Cox, N. J., et al. (2016). Metaxcan: summary statistics based gene-level association method infers accurate predixcan results. BioRxiv, page 045260.

Carbonetto, P. and Stephens, M. (2013). Integrated enrichment analysis of variants and pathways in genome-wide association studies indicates central role for il-2 signaling genes in type 1 diabetes, and cytokine signaling genes in crohn’s disease. PLoS genetics, 9(10), e1003770.

Chang, Y., Glass, K., Liu, Y.-Y., Silverman, E. K., Crapo, J. D., Tal-Singer, R., Bowler, R., Dy, J., Cho, M., and Castaldi, P. (2016). Copd subtypes identified by network-based clustering of blood gene expression. Genomics, 107(2), 51–58.

Efron, B. (2012). Large-scale inference: empirical Bayes methods for estimation, testing, and prediction, volume 1. Cambridge University Press.

Elovainio, M., Taipale, T., Seppµlµ, I., Mononen, N., Raitoharju, E., Jokela, M., Pulkki-R°aback, L., Illig, T., Waldenberger, M., Hakulinen, C., et al. (2015). Activated immune–inflammatory pathways are associated with long-standing depressive symptoms: evidence from gene-set enrichment analyses in the young finns study. Journal of psychiatric research, 71, 120–125.

Gamazon, E. R., Wheeler, H. E., Shah, K. P., Mozaffari, S. V., Aquino-Michaels, K., Carroll, R. J., Eyler, A. E., Denny, J. C., Nicolae, D. L., Cox, N. J., et al. (2015). A gene-based association method for mapping traits using reference transcriptome data. Nature genetics, 47(9), 1091.

Gusev, A., Mancuso, N., Won, H., Kousi, M., Finucane, H. K., Reshef, Y., Song, L., Safi, A., McCarroll, S., Neale, B. M., et al. (2018). Transcriptome-wide association study of schizophrenia and chromatin activity yields mechanistic disease insights. Nature genetics, 50(4), 538.

Guttman, M., Amit, I., Garber, M., French, C., Lin, M. F., Feldser, D., Huarte, M., Zuk, O., Carey, B. W., Cassady, J. P., et al. (2009). Chromatin signature reveals over a thousand highly conserved large non-coding rnas in mammals. Nature, 458(7235), 223.

Hass, J., Walton, E., Wright, C., Beyer, A., Scholz, M., Turner, J., Liu, J., Smolka, M. N., Roessner, V., Sponheim, S. R., et al. (2015). Associations between dna methylation and schizophrenia-related intermediate phenotypes—a gene set enrichment analysis. Progress in Neuro-Psychopharmacology and Biological Psychiatry, 59, 31–39.

Keshava Prasad, T., Goel, R., Kandasamy, K., Keerthikumar, S., Kumar, S., Mathivanan, S., Telikicherla, D., Raju, R., Shafreen, B., Venugopal, A., et al. (2008). Human protein reference database—2009 update. Nucleic acids research, 37(Suppl 1), D767–D772.

Love, M. I., Huber, W., and Anders, S. (2014). Moderated estimation of fold change and dispersion for rna-seq data with deseq2. Genome biology, 15(12), 550.

Maruschke, M., Hakenberg, O. W., Koczan, D., Zimmermann, W., Stief, C. G., and Buchner, A. (2014). Expression profiling of metastatic renal cell carcinoma using gene set enrichment analysis. International Journal of Urology, 21(1), 46–51.

Mootha, V. K., Lindgren, C. M., Eriksson, K.-F., Subramanian, A., Sihag, S., Lehar, J., Puigserver, P., Carlsson, E., Ridderstr°ale, M., Laurila, E., et al. (2003). Pgc-1*α*-responsive genes involved in oxidative phosphorylation are coordinately downregulated in human diabetes. Nature genetics, 34(3), 267.

Moyerbrailean, G. A., Richards, A. L., Kurtz, D., Kalita, C. A., Davis, G. O., Harvey, C. T., Alazizi, A., Watza, D., Sorokin, Y., Hauff, N., et al. (2016). High-throughput allele-specific expression across 250 environmental conditions. Genome research, pages gr–209759.

Richiardi, J., Altmann, A., Milazzo, A.-C., Chang, C., Chakravarty, M. M., Banaschewski, T., Barker, G. J., Bokde, A. L., Bromberg, U., Büchel, C., et al. (2015). Correlated gene expression supports synchronous activity in brain networks. Science, 348(6240), 1241–1244.

Schaub, F. X., Dhankani, V., Berger, A. C., Trivedi, M., Richardson, A. B., Shaw, R., Zhao, W., Zhang, X., Ventura, A., Liu, Y., et al. (2018). Pan-cancer alterations of the myc oncogene and its proximal network across the cancer genome atlas. Cell systems, 6(3), 282–300.

Segrè, A. V., Groop, L., Mootha, V. K., Daly, M. J., Altshuler, D., Consortium, D., Investigators, M., et al. (2010). Common inherited variation in mitochondrial genes is not enriched for associations with type 2 diabetes or related glycemic traits. PLoS genetics, 6(8), e1001058.

Shalem, O., Sanjana, N. E., Hartenian, E., Shi, X., Scott, D. A., Mikkelsen, T. S., Heckl, D., Ebert, B. L., Root, D. E., Doench, J. G., et al. (2014). Genome-scale crispr-cas9 knockout screening in human cells. Science, 343(6166), 84–87.

Speliotes, E. K., Willer, C. J., Berndt, S. I., Monda, K. L., Thorleifsson, G., Jackson, A. U., Allen, H. L., Lindgren, C. M., Luan, J., Mµgi, R., et al. (2010). Association analyses of 249,796 individuals reveal 18 new loci associated with body mass index. Nature genetics, 42(11), 937.

Stephens, M. (2016). False discovery rates: a new deal. Biostatistics, 18(2), 275–294.

Storey, J. D. et al. (2003). The positive false discovery rate: a bayesian interpretation and the q-value. The Annals of Statistics, 31(6), 2013–2035.

Subramanian, A., Tamayo, P., Mootha, V. K., Mukherjee, S., Ebert, B. L., Gillette, M. A., Paulovich, A., Pomeroy, S. L., Golub, T. R., Lander, E. S., et al. (2005). Gene set enrichment analysis: a knowledge-based approach for interpreting genome-wide expression profiles. Proceedings of the National Academy of Sciences, 102(43), 15545–15550.

Walter, N. D., Dolganov, G. M., Garcia, B. J., Worodria, W., Andama, A., Musisi, E., Ayakaka, I., Van, T. T., Voskuil, M. I., De Jong, B. C., et al. (2015). Transcriptional adaptation of drug-tolerant mycobacterium tuberculosis during treatment of human tuberculosis. The Journal of infectious diseases, 212(6), 990–998.

Willer, C. J., Schmidt, E. M., Sengupta, S., Peloso, G. M., Gustafsson, S., Kanoni, S., Ganna, A., Chen, J., Buchkovich, M. L., Mora, S., et al. (2013). Discovery and refinement of loci associated with lipid levels. Nature genetics, 45(11), 1274.

Zhu, Z., Zhang, F., Hu, H., Bakshi, A., Robinson, M. R., Powell, J. E., Montgomery, G. W., Goddard, M. E., Wray, N. R., Visscher, P. M., et al. (2016). Integration of summary data from gwas and eqtl studies predicts complex trait gene targets. Nature genetics, 48(5), 481.

