## Supplemental Figures for "BAGSE: a Bayesian hierarchical model approach for gene set enrichment analysis" (from another version of this preprint: version 1, 2019-06-06)

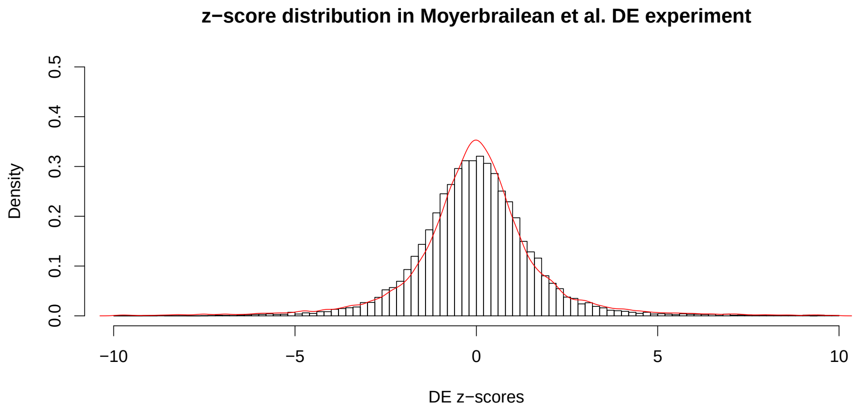


Supplementary Figure 1: Comparison of empirical $z$-score distribution with a $t$-distribution with 10 degree of freedom. The histogram represents the empirical $z$-score distribution from Moyerbrailean et al. (2017). The read line represents the theoretical density function of the $t$-distribution with 10 degree of freedom.

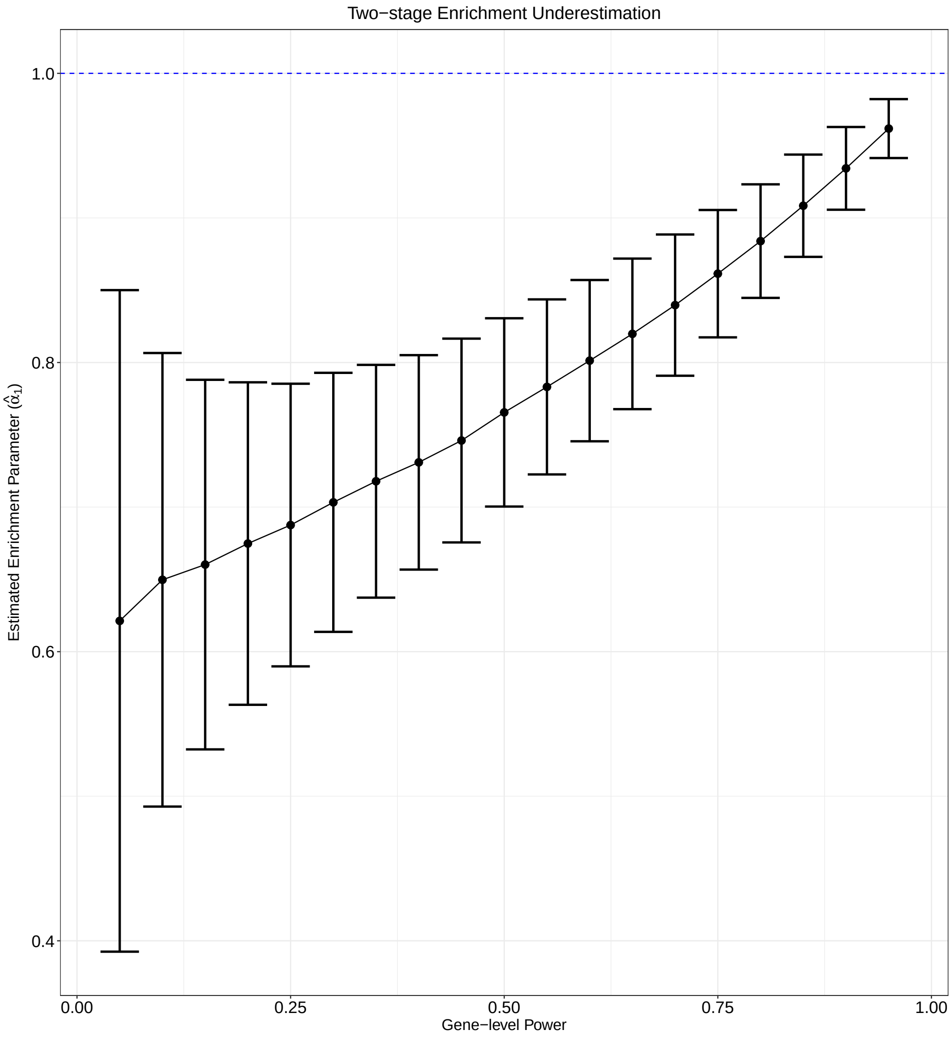


Supplementary Figure 2: A demonstration of the underestimation of the enrichment log odds ratio as a function of power in gene-level tests for the two-stage approach. In this example, the true enrichment parameter value $\alpha_1$ is 1.

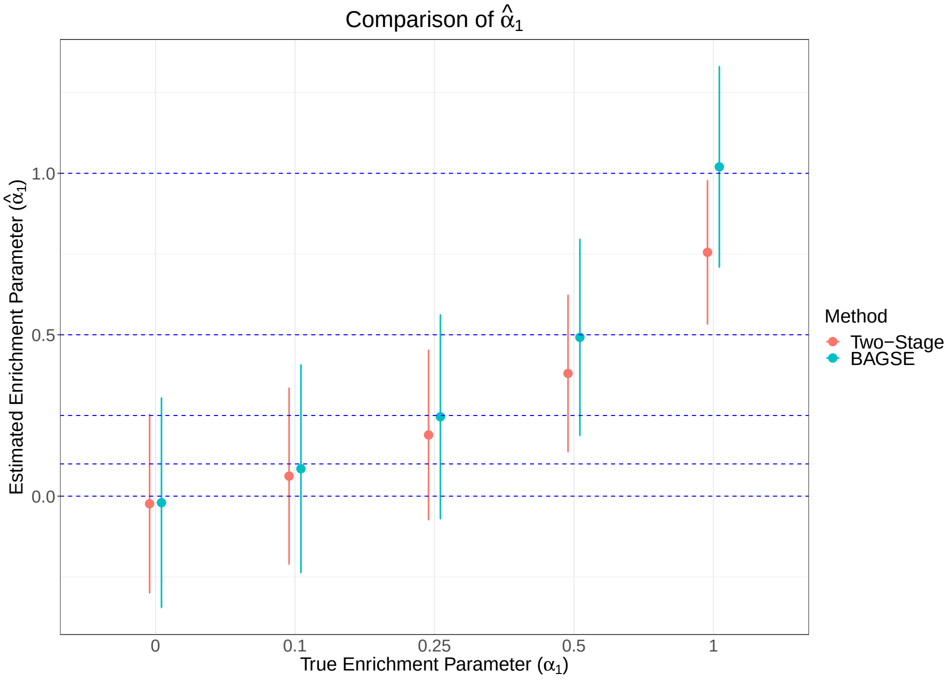


Supplementary Figure 3: A comparison of the two-stage approach and BAGSE in terms of estimation of $\alpha_1$, when 1% of genes are annotated.

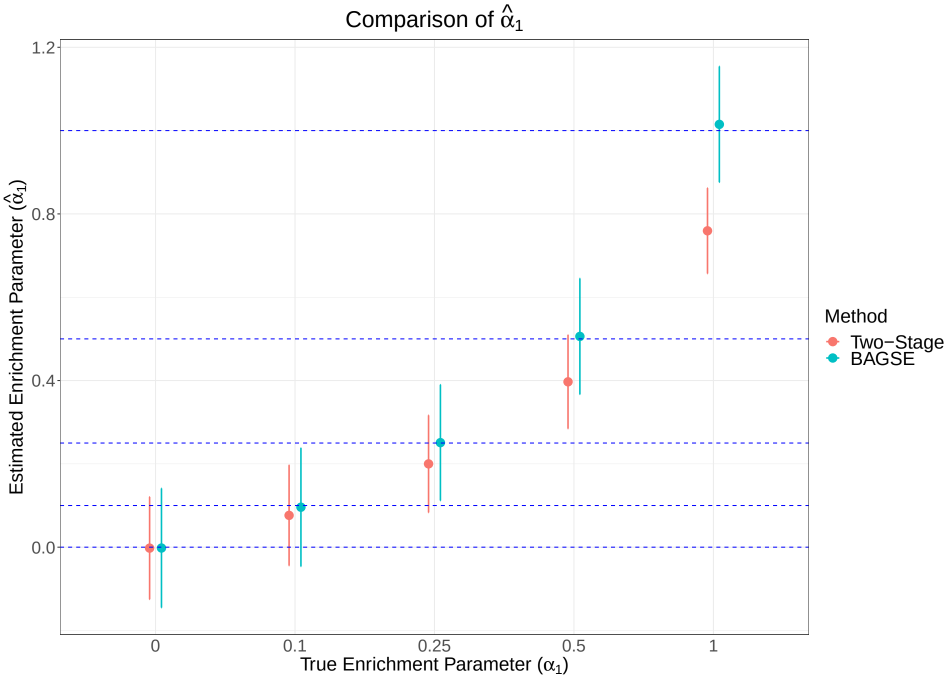


Supplementary Figure 4: A comparison of the two-stage approach and BAGSE in terms of estimation of $\alpha_1$, when 5% of genes are annotated.

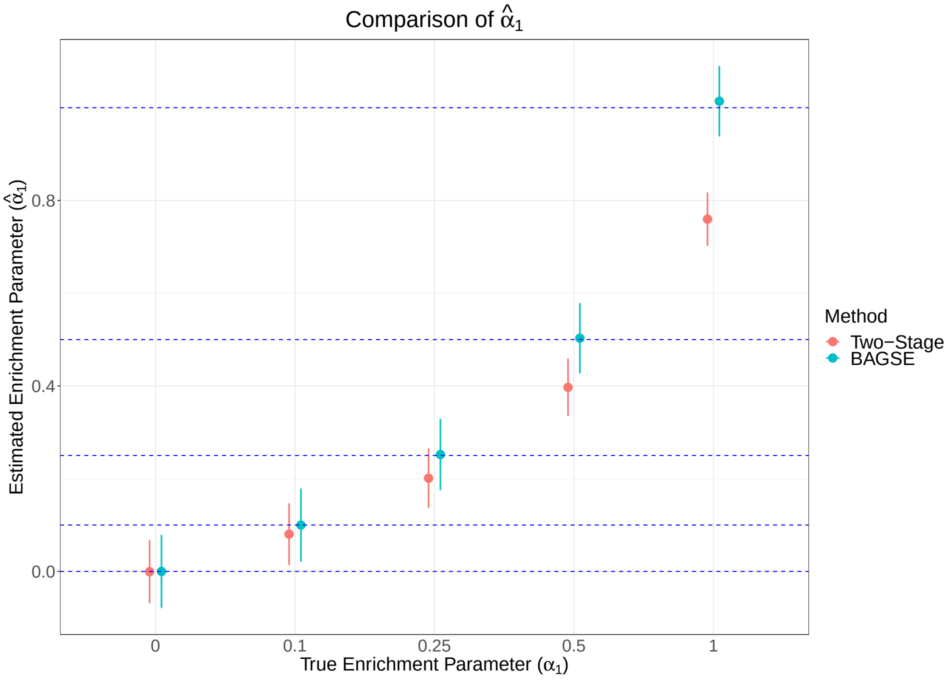


Supplementary Figure 5: A comparison of the two-stage approach and BAGSE in terms of estimation of $\alpha_1$, when 20% of genes are annotated.

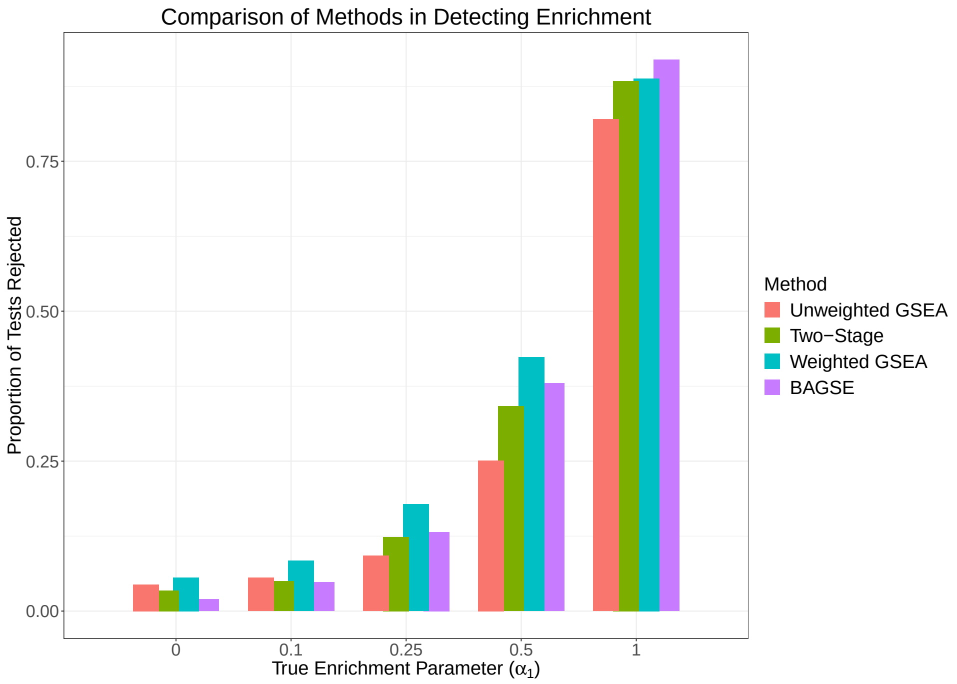


Supplementary Figure 6: A comparison of the proportion of tests for detecting enrichment between the unweighted GSEA, two-stage approach, weighted GSEA, and BAGSE, when 1% of genes are annotated

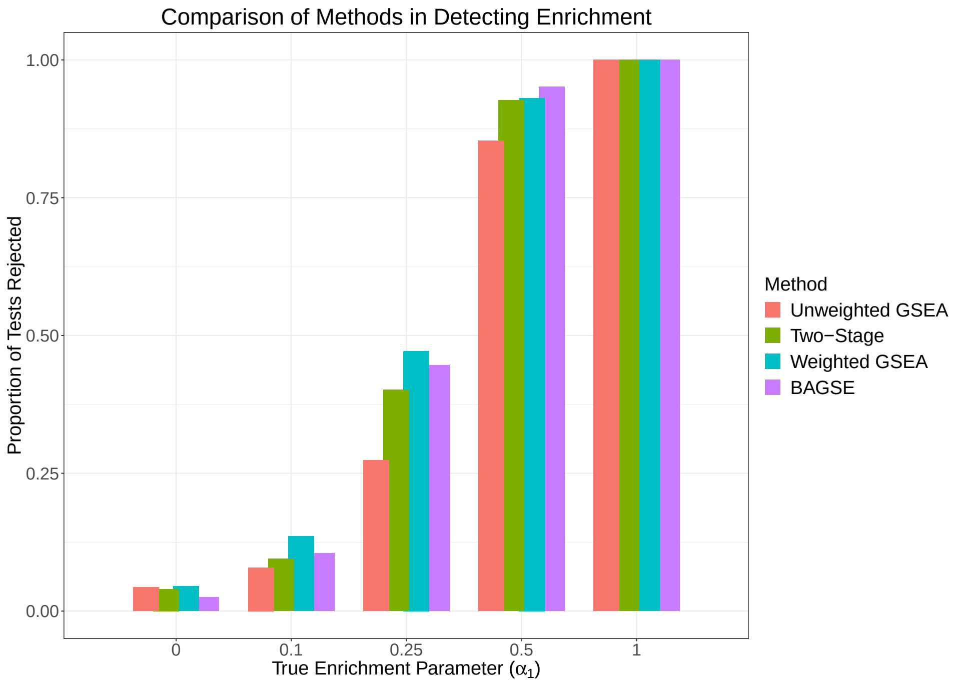


Supplementary Figure 7: A comparison of the proportion of tests for detecting enrichment between the unweighted GSEA, two-stage approach, weighted GSEA, and BAGSE, when 5% of genes are annotated

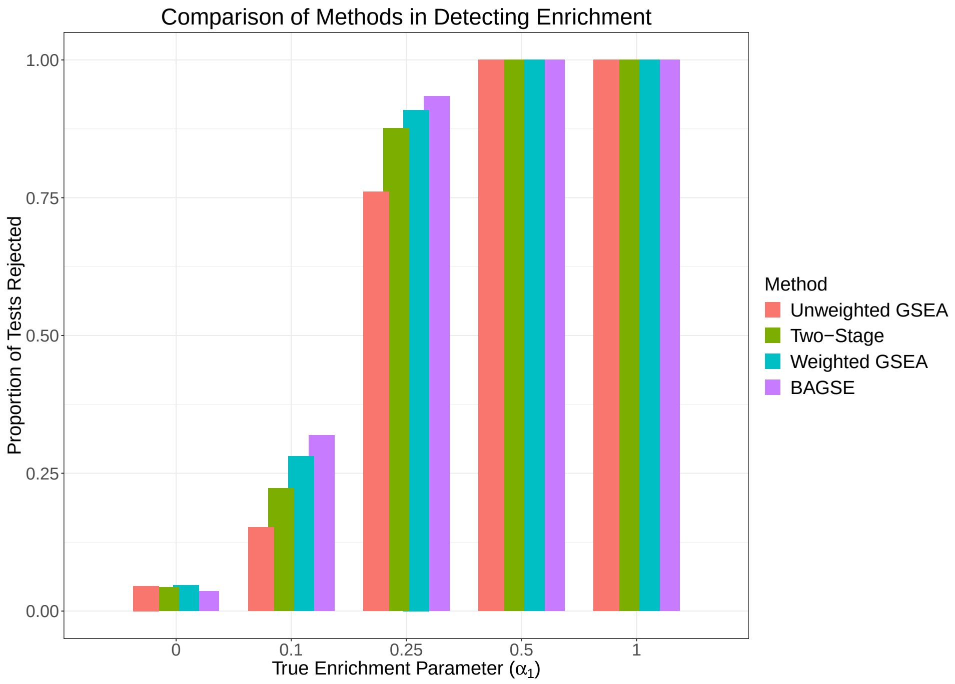


Supplementary Figure 8: A comparison of the proportion of tests for detecting enrichment between the unweighted GSEA, two-stage approach, weighted GSEA, and BAGSE, when 20% of genes are annotated
