## Supplemental Table 1 for "BAGSE: a Bayesian hierarchical model approach for gene set enrichment analysis" (from another version of this preprint: version 1, 2019-06-06)

Supplementary Table 1:

The results of all our simulation studies comparing the ability to quantify enrichment level between BAGSE and the two-stage approach. The table below contains these enrichment level estimates for BAGSE and the two-stage approach for each parameter set.

| Proportion of Genes Annotated | Number of Genes Annotated | Method | True Enrichment Parameter | | | | | Truly Associated Annotated genes | | | | |
|---|---|---|---|---|---|---|---|---|---|---|---|---|
| | | | **0** | **0.1** | **0.25** | **0.5** | **1** | **0** | **0.1** | **0.25** | **0.5** | **1** |
| **0.01** | **100** | BAGSE | -0.02 | 0.085 | 0.246 | 0.492 | 1.02 | 27 | 27 | 31 | 39 | 47 |
| | | Two-stage | -0.02 | 0.062 | 0.19 | 0.38 | 0.755 | | | | | |
| **0.05** | **500** | BAGSE | -0.002 | 0.096 | 0.251 | 0.506 | 1.015 | 127 | 134 | 153 | 185 | 235 |
| | | Two-Stage | -0.002 | 0.076 | 0.2 | 0.397 | 0.759 | | | | | |
| **0.1** | **1000** | BAGSE | -0.001 | 0.098 | 0.249 | 0.506 | 1.015 | 267 | 283 | 314 | 377 | 493 |
| | | Two-Stage | -0.001 | 0.079 | 0.199 | 0.397 | 0.76 | | | | | |
| **0.2** | **2000** | BAGSE | 0 | 0.1 | 0.252 | 0.503 | 1.014 | 532 | 572 | 622 | 738 | 990 |
| | | Two-Stage | 0 | 0.08 | 0.201 | 0.397 | 0.76 | | | | | |
