## Supplemental Table 2 for "BAGSE: a Bayesian hierarchical model approach for gene set enrichment analysis" (from another version of this preprint: version 1, 2019-06-06)

### Supplementary Table 2:

The results of all our simulation studies comparing the power to detect enrichment between all four examined methods. The table below contains these estimates of power and rate of type I error control for each parameter set.

| Proportion of Genes Annotated | Number of Genes Annotated | Truly Associated and Annotated genes | | | | | Method | True Enrichment Parameter | | | | |
|---|---|---|---|---|---|---|---|---|---|---|---|---|
| | | $\alpha_1$= 0 | $\alpha_1$= 0.1 | $\alpha_1$= 0.25 | $\alpha_1$= 0.5 | $\alpha_1$= 1 | | **0 (False Positives)** | **0.1** | **0.25** | **0.5** | **1** |
| **0.01** | **100** | 27 | 27 | 31 | 39 | 47 | BAGSE | 0.02 | 0.048 | 0.132 | 0.38 | 0.919 |
| | | | | | | | Weighted GSEA | 0.056 | 0.084 | 0.178 | 0.423 | 0.888 |
| | | | | | | | Two-stage | 0.034 | 0.049 | 0.124 | 0.342 | 0.884 |
| | | | | | | | Unweighted GSEA | 0.044 | 0.055 | 0.092 | 0.251 | 0.82 |
| **0.05** | **500** | 127 | 134 | 153 | 185 | 235 | BAGSE | 0.025 | 0.11 | 0.446 | 0.951 | 1 |
| | | | | | | | Weighted GSEA | 0.045 | 0.136 | 0.472 | 0.931 | 1 |
| | | | | | | | Two-stage | 0.04 | 0.095 | 0.402 | 0.927 | 1 |
| | | | | | | | Unweighted GSEA | 0.043 | 0.079 | 0.274 | 0.854 | 1 |
| **0.1** | **1000** | 267 | 283 | 314 | 377 | 493 | BAGSE | 0.03 | 0.178 | 0.717 | 0.999 | 1 |
| | | | | | | | Weighted GSEA | 0.056 | 0.214 | 0.707 | 0.997 | 1 |
| | | | | | | | Two-stage | 0.047 | 0.147 | 0.637 | 0.997 | 1 |
| | | | | | | | Unweighted GSEA | 0.046 | 0.11 | 0.484 | 0.991 | 1 |
| **0.2** | **2000** | 532 | 572 | 622 | 738 | 990 | BAGSE | 0.036 | 0.319 | 0.934 | 1 | 1 |
| | | | | | | | Weighted GSEA | 0.047 | 0.281 | 0.909 | 1 | 1 |
| | | | | | | | Two-stage | 0.043 | 0.223 | 0.876 | 1 | 1 |
| | | | | | | | Unweighted GSEA | 0.046 | 0.152 | 0.761 | 1 | 1 |
