## Supplemental Table 3 for "BAGSE: a Bayesian hierarchical model approach for gene set enrichment analysis" (from another version of this preprint: version 1, 2019-06-06)

### Supplementary Table 3:

The BAGSE enrichment estimate and 95% Confidence Interval for all pathways within the Immune Response category in KEGG. The quantified enrichment estimate's interpretation as a log odds ratio leads to easily comparable quantities. Please note that certain of the Immune Response pathways did not have any genes in our data, and were therefore not included.

| Pathway | Number of Genes | BAGSE Enrichment Estimate | 95% C.I |
|---|---|---|---|
| Intestinal Immune Network for IgA Production | 38 | 9.34 | (5.38, 13.29) |
| Hematopoietic Cell Lineage | 49 | 2.69 | (1.33, 4.04) |
| Cytosolic DNA-sensing Pathway | 36 | 2.41 | (0.73, 4.08) |
| B Cell Receptor Signaling Pathway | 73 | 1.84 | (0.65, 3.04) |
| RIG-I-like Receptor Signaling Pathway | 54 | 1.69 | (0.59, 2.79) |
| Toll-like Receptor Signaling Pathway | 77 | 1.53 | (0.54, 2.51) |
| Antigen Processing and Presentation | 47 | 1.41 | (0.10, 2.72) |
| Natural Killer Cell Mediated Cytotoxicity | 87 | 1.26 | (0.32, 2.20) |
| Fc Gamma R-mediated Phagocytosis | 84 | 1.05 | (0.05, 2.06) |
| T Cell Receptor Signaling Pathway | 96 | 1.04 | (0.11, 1.97) |
| Leukocyte Transendothelial Migration | 90 | 0.97 | (0.07, 1.87) |
| Chemokine Signaling Pathway | 132 | 0.8 | (0.03, 1.57) |
| NOD-like Receptor Signaling Pathway | 48 | 0.51 | (-1.23, 2.25) |
| Fc Epsilon RI Signaling Pathway | 66 | 0.37 | (-5.13, 5.87) |
| Complement and Coagulation Cascades | 27 | -0.25 | (-2.04, 1.55) |
